# Comparing adult hippocampal neurogenesis across species: translating time to predict the tempo in humans

**DOI:** 10.1101/404202

**Authors:** Christine J. Charvet, Barbara L Finlay

**Affiliations:** Department of Psychology, Delaware State University, Dover, DE, USA; Laboratory of Behavioral and Evolutionary Neuroscience, Department of Psychology, Cornell University, Ithaca, NY, USA.; Laboratory of Behavioral and Evolutionary Neuroscience, Department of Psychology, Cornell University, Ithaca, NY, USA

**Keywords:** hippocampus, neurogenesis, adult, human, rodent, monkey, Ki67, allometry

## Abstract

Comparison of neurodevelopmental sequences between species whose initial period of brain organization may vary from one hundred days to one thousand days, and whose progress is intrinsically nonlinear presents large challenges in normalization. Comparing adult timelines when lifespans stretch from one year to seventy-five, when underlying cellular mechanisms under scrutiny do not scale similarly, presents challenges to simple detection and comparison. The question of adult hippocampal neurogenesis has generated numerous controversies regarding its simple presence or absence in humans versus rodents, whether it is best described as the tail of a distribution centered on early neural development, or is several distinct processes. In addition, adult neurogenesis may have substantially changed in evolutionary time in different taxonomic groups. Here we extend and adapt a model of the cross-species transformation of early neurodevelopmental events which presently reaches up to the equivalent of the third human postnatal year for 18 mammalian species (www.translatingtime.net) to address questions relevant to hippocampal neurogenesis, which permit extending the database to adolescence or perhaps to the whole lifespan. We acquired quantitative data delimiting the envelope of hippocampal neurogenesis from cell cycle markers (i.e., Ki67, DCX) and RNA sequencing data for two primates (macaque, humans) and two rodents (rat, mouse). To improve species coverage in primates, we gathered the same data from marmosets (*Callithrix jacchus*), but additionally gathered data on a number of developmental milestones to find equivalent developmental time points between marmosets and other species. When all species are so modeled, and represented in a common time frame, the envelopes of hippocampal neurogenesis are essentially superimposable. Early developmental events involving the olfactory and limbic system start and conclude possibly slightly early in primates than rodents, and we find a comparable early conclusion of primate hippocampal neurogenesis (as assessed by the relative number of Ki67 cells) suggesting a plateau to low levels at approximately 2 years of age in humans. Marmosets show equivalent patterns within neurodevelopment, but unlike macaque and humans may have wholesale delay in the initiation of neurodevelopment processes previously observed in some precocial mammals such as the guinea pig and multiple large ungulates.

## 1.0 Introduction

The following paper, a contribution to the collection “Adult Neurogenesis: beyond rats and mice”, is a hybrid of two components. At its core is an empirical contribution to the literature on hippocampal neurogenesis, comparing late neurogenesis in two rodents and three primates, using evidence from cell cycle markers. We have informed that analysis with the “Translating Time” project (www.translatingtime.net), where we have gathered evidence about the relative progress of neurodevelopmental events from the first birthdays of mature neurons until increasingly later ages across 18 mammalian species. We will argue that any claim that onset, offset or duration of a developmental process, or an adult brain feature produced by such a process, is “unique”, or even “specialized” in humans or any other species or taxonomic group is absolutely dependent on a proper allometric comparison, such as made possible by the “Translating time” modeling work, or other similar analyses. A comparison of a developmental feature of the brain of a particular rodent to particular primate species is not such an analysis, and will systematically mislead researchers.

The second component is a discussion of the problems and opportunities of developmental allometric analyses across mammals, which we present in this introduction. We include a review and exposition of basic allometric claims and procedures as they apply to brain mass and developmental duration in general, as well as the progress of neurogenesis targeted in this paper. We will describe some of the quantitative misunderstandings that typically arise from moving between the exponential functions used in allometric analyses, and the linear functions used in basic measurements of cell number and volume in developmental cell biology. The immediately following expanded introduction concerns the motivation, history, and methodology necessary to understand methods of analysis in developmental allometry.

### 1.1 The purpose and methodology of allometric comparison

#### 1.1.1 Prior work on developmental allometry

In order to limit the need to contrast statistical methodologies of successive papers within the text, the following description of species, neural structures, developmental span and mathematical models employed in this research project follows, including immediately relevant work of several other laboratories. First, the current database for the “Translating Time” model with tables of species, structures and sources, a description of the current model, and a utility to translate or predict a developmental equivalent day between any two species in the model can be found at www.translatingtime.org (Clancy et al., 2007). The initial comparison of neurogenetic schedules in rhesus monkey, cat, four rodents and a marsupial, extending from onset of neurogenesis to approximately birth in the monkey, using principal components analysis, is described in Finlay and Darlington (1995) and an extended discussion of statistical considerations, principally phylogenetic covariation can be found in Finlay et al. (2001). Darlington et al. (1999) and Clancy et al. (2000) bring the number of species to 9 eutherian (placental) mammals including humans and 6 metatherians (marsupials), principally using regression analyses. Clancy et al. (2001) extend the neurodevelopmental events past neurogenesis to include synaptogenesis, cell death, ocular dominance columns and the like, using regression and the general linear model (see also Clancy et al., 2008). The relationship of individual variability to between-species variability is discussed in Finlay et al. (2011), and specifically in humans in Charvet et al. (2013). The current iterative model deriving the “event scale” of maturation developed in Workman et al. (2013) brings the number of mammalian species to 18, the number of developmental events to 271, including myelination, volume change, and early behavioral events, extending to human-equivalent of the third postnatal year. Particularly relevant to the present paper, patterns in the neural maturation of altricial versus precocial species are contrasted. A demonstration of the problems arising from a failure to account for allometric concerns can be found in “Human exceptionalism” (Finlay and Workman, 2013). Early behavioral development and related neuroplasticity are integrated with translating time in Finlay and Uchiyama (2017), and finally, evolution of life histories, including events like weaning and menopause in Hawkes and Finlay (2018). Readers are directed to the early work of Passingham (1985), and Garwicz et al. (2009), who use similar methods to examine early independent ambulation, as well as that of Halley’s studies of the growth of initial primordia and brain across a wide range of mammals (Halley, 2016, 2017). More recently, the advent of single cell RNA sequencing provides an exciting opportunity to investigate developmental trajectories of neural subpopulations across species (Habib et al., 2017; Iacono et al., 2017; Fan et al., 2018; Zhong et al., 2018). We here broaden the maturational range of neurodevelopmental ages of studies in our database to capture late stages of hippocampal neurogenesis across species.

#### 1.1.2 Allometry of brain and brain parts

The general form of scaling of neural mass or neuron numbers in any brain region compared to the whole brain, has been studied for many years (Jerison, 1973; Gould, 1975; Fleagle, 1985). Overall consensus exists about general features of brain and body scaling, though subject to the normal continuing debate about optimal ways to quantify statistical variation in large and complex datasets (Finlay et al., 2001; Freckleton et al., 2002). We will take the particular example of cross-species comparisons of the volume and number of neurons in the neocortex, and particularly the frontal cortex (the allometric study of the brain), to introduce the related and less familiar topic of scaling of developmental duration across species, which we term developmental allometry.

If scaling of neocortical volume (or “isocortex”) is the focus for consideration, the fact that the human brain has a disproportionately large cortex compared to primates and most other mammals is quite “obvious” – for example, the human cortex comprises over 80% of its total brain mass, compared to around 20% in shrews and rodents (Finlay and Darlington, 1995). The correct empirical observation of the apparently disproportionate size of the cortex along with its persistent misinterpretation is the prototypical example of a problem we will call “human exceptionalism” (Finlay and Workman, 2013). The disproportionate volume of the human neocortex suggested to multiple researchers alike -- anthropologists, embryologists, neuroscientists and psychologists -- that it must be the result of special selection compared to the rest of the brain. Since the cortex was thus thought to be the subject of selection within the brain, every cognitive alteration or adaptation in evidence in humans has typically been typically credited to its superior computational prowess. But it’s not necessarily so. Although we have an unusually large brain, our cortex is the size it should be for a brain of our absolute size when cross-species cortex volume or cell numbers are represented on logarithmic scales (Jerison, 1973; Hofman, 1989; Finlay and Darlington, 1995; Kaas and Herculano-Houzel, 2017).

#### 1.1.3 Linear scales, logarithmic scales and the allometric equation

A “proper” comparison of variations across species of different sizes and developmental durations requires care (this section is abridged from Hawkes and Finlay, 2018 to which the reader is directed for a more extensive discussion). Even with “all else equal” in such factors as a species’ niche, number of brain components, sex and age, still, the laws of geometry, and of physics and chemistry, impose lawful changes in both form and process with increase in brain mass. The intrinsic geometry of physical relationships results in variable allometric relationships (e.g., doubling the volume of a sphere only increases its radius by 1.26 times). After such geometric constants are understood, any two structures or processes changing in size or duration across species could show nonlinear scaling relationships, scale linearly, or might show no predictable scaling, depending on the mechanisms or functions that are relevant. For example, the divisions, doubling and redoubling of stem cell pools are best described by nonlinear equations. Some features change linearly: for example, if multiplied by the appropriate constant, cross-sectional diagrams of mice and rat eyes are superimposable even though the rat’s eye is twice as big, as both are solving a linear optical problem with the same materials (Remtulla and Hallet, 1985). Some features do not scale at all with brain mass (considering mammals only here), such as the diameter of the cell bodies of neurons, the time to complete the first generation of a mature neuron, or the duration of action potentials. Such variable geometrical and biological scaling relations can coexist for different aspects of the same structure. Finally, datasets of interest often have underlying geometries that can mislead graphical comparisons. Consider a typical Mercator projection of the earth’s landmasses, where the continents of Africa and Greenland appear approximately equal in size, but when measured in its correct spherical coordinates, Africa is more than 10 times larger than Greenland.

Allometries are conventionally represented as scaling relationships. If the relationship between two features that correlate with each other in size, say ‘x’ and ‘y,’ is represented as y=kx^a^ where ‘k’ is some constant, and the exponent ‘a’ represents the rate at which ‘y’ changes with respect to a change in ‘x.’ If exponent ‘a’ is more or less than 1 then a change in y is associated with a geometrical change in ‘x.’ Such geometrical or exponential relationships can be plotted and visualized as linear ones by logarithmic transformation: log y = *a*log *x* + log *k*. In such log plots, the exponent ‘a’ now appears as the slope of the increase in y with respect to x. Using this representation of cortex mass relative to the whole brain is represented on a logarithmic scale, it is clear that the human neocortex is exactly the size it “should” be (Figure 1). The human brain is absolutely large compared to other primates, but given this large brain size, each part falls onto its “expected” position, from hindbrain to cortex (Hofman, 1989). The cortex has “positive allometry” with respect to the rest of the brain, its slope greater than one, which is the “linear scaling reference” of Figure 1. Inevitably, therefore, with different brain components each increasing in mass at different rates, larger mammalian brains become “disproportionately” composed of cortex. The exact exponent of cortical positive allometry might vary with whether neurons, all cells, surface area or volume is measured, and shows some taxon-specific differences, but none reduce the positive exponent to one or less (a sampling of a large literature: Jerison, 1973; 1989; Hofman, 1989; Reep et al., 2007; Herculano-Houzel et al., 2017; Charvet et al., 2013).

**Figure 1.**
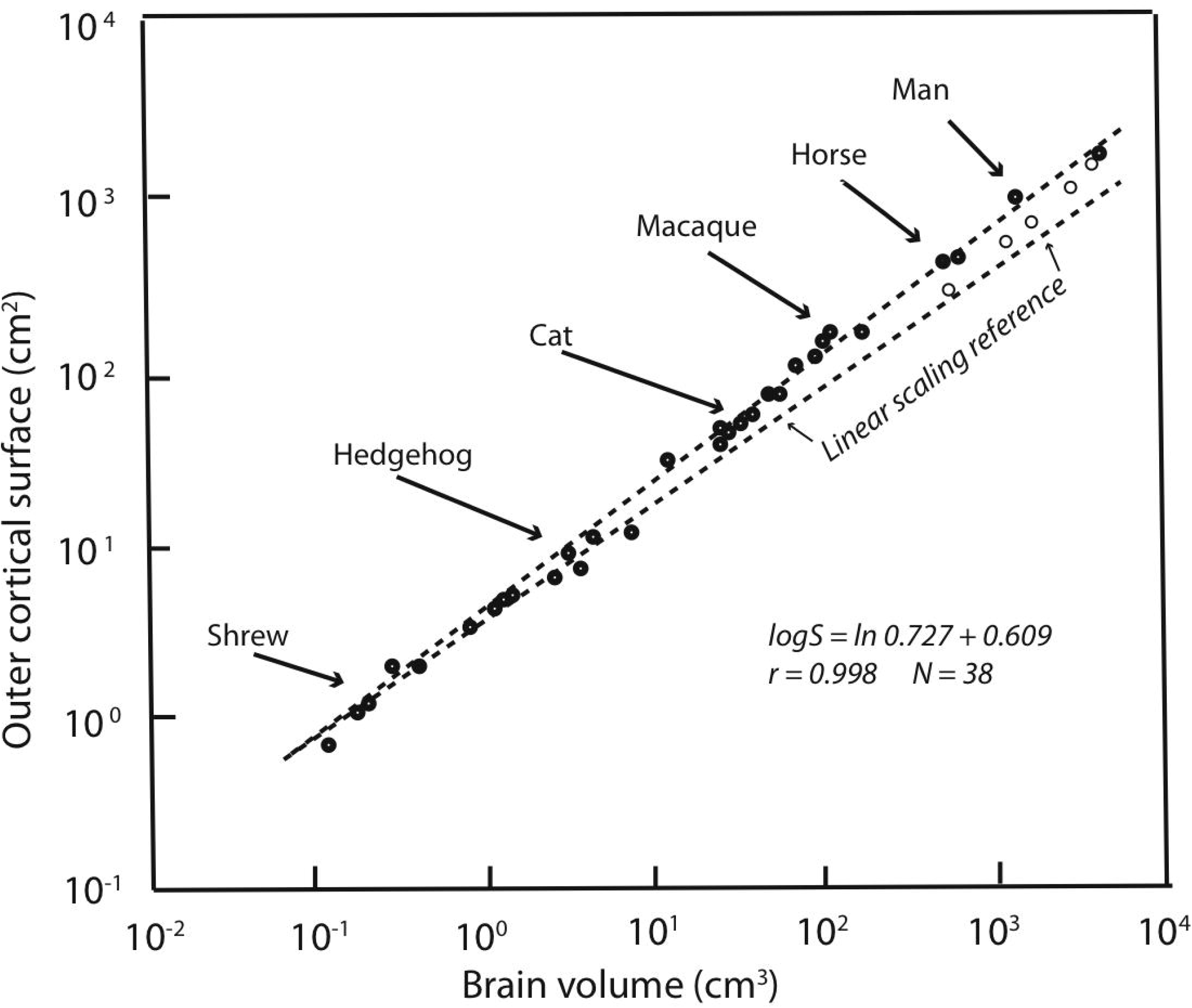
Redrawn from Figure 2 in Hofman, 1989. Outer cortical surface area is plotted as a function of brain volume on a logarithmic scale. The slope of the standard major axis is 0.727 +/− 0.009. The dashed line represents the scaling of the cortical surface area according to a two-thirds power relation (the necessary geometric similarity of plotting an area against a volume if both increase linearly). Dolphins and whales are indicated by circles.

Because of the regular, predictable relationships of the relative sizes of brain parts at all absolute brain volumes, lacking other information, our large cortex cannot be attributed to special selection for that feature, as it comes “for free” with selection on the whole brain, or in fact, could arise by leverage by selection on any part of the brain (Finlay and Darlington, 1995). It is interesting, to be sure, that over evolutionary time that the cortex, and the cerebellum are the two brain regions where disproportionate neuron number, volume and energy consumption are routinely allocated (Finlay et al., 2011). Comparison of relative cortical and cerebellar volume between any two mammals of different brain size will reveal this feature, not only comparison of the human brain with all others. The most telling evidence is that those several mammalian brains which are absolutely larger in mass than the human brain, including several cetaceans and ungulates, continue the allometric equation of the cortex, so that they have proportionately even more cortex than humans do (Figure 1).

#### 1.1.4 The evolutionary question at issue: the case of the prefrontal cortex

Questions involving allometric scaling are in no way historical debates as a similar controversy is ongoing about whether a specific region of cortex, the prefrontal cortex, is “allometrically unexpected” in humans (Sherwood and Smaers, 2013). Just as the cortex has a particular exponent of enlargement with respect to the rest of the brain, every cortical area (e.g., prefrontal, primary visual) has its own exponent (or slope in the log-transformed equation) showing its change in relative volume compared to overall cortex volume. Both the prefrontal and parietal cortex regions have an exponent that is larger than the cortex’s overall exponent, showing a positive allometry (Jerison, 1997). The issue under debate is whether the frontal cortex in humans is larger still than would be expected from its already high positive allometry (Barton and Venditti, 2013; Chaplin, Yu, Soares, Gattass, and Rosa, 2013; Passingham and Smaers, 2014; Semendeferi, Lu, Schenker, and Damasio, 2002). As before, however, when we discussed preferential allocation of “excess” neural mass for cortex and cerebellum versus the rest of the brain, it is interesting that it is frontal and parietal cortex that are preferentially enlarged in the cortical sheet when brains increase in volume across mammals.

Why should these researchers care about this issue? If researchers claim a region’s volume is “allometrically unexpected” in humans, they are claiming that it must have been the target of selection, typically because of special importance of the function ascribed to that brain region in that species. In the case of the frontal cortex, the cognitive features usually evoked are cognitive control, the ability to choose reasonable behavioral solutions from competing possibilities, or to evaluate choices with respect to goals distant in space or time. Thus, the claim that the frontal cortex is allometrically unexpected in humans is a claim that humans have been selected on a behavioral feature like cognitive control, which in turn is improved with the relative volume of frontal cortex. Structures that change their volume according to regular, cross-species allometric rules, however, even if they look disproportionate on a linear scale, require no special explanation. If the entire brain has been under special selection for larger size in any species, every single change in the proportionality of its parts is generated by its change in size. We’ll make no ruling on this claim, except to note that the deviation in human frontal cortex volume, if it exists, is small enough to make it susceptible to relatively minor differences in methodology between research groups.

It remains interesting and important that brains enlarge in particular ways, and that predictable patterns of reorganization, both behavioral and computational, are associated with cortical enlargement (Finlay and Uchiyama, 2015). Mammals with large brains are certain to show evidence of a disproportionate contribution of frontal cortex (Passingham and Smaers, 2014). Allometric regularities in structural scaling, whether in the cortex, or in the hippocampus we will soon be discussing, require that we investigate coordinated mechanisms *outside* the structures of interest, and should make us skeptical of causal accounts that depend on selection on hypothesized special adaptations of the particular species of animal.

An important mechanism of volumes and neuron number coordination in several cases studied so far appears to be the coordinated control of duration of neurogenesis, as applied to every part of mammalian brains (e.g. Cahalane et al., 2014; Charvet and Finlay, 2014; Dyer et al., 2009; Finlay and Darlington, 1995). As the duration of hippocampal neurogenesis is the subject of the empirical component of this paper, we will now turn to issues in the allometry of development.

### 1.2 The allometry of developmental duration: basic requirements

#### 1.2.1 The need for data from multiple species: why attempts to “norm” measurements between only two species will be ineffective

The formal properties of “allometrically expected” changes in mass also apply to translations of developmental time from one species to another. The appropriate coordinate system to represent time translations will depend on the data to be represented, and the representation desired. The relationship of developmental timing between species cannot be presumed to be best represented on a linear scale. In order to fairly compare developmental durations between animals, enough data must be collected from a number of relevant species to support generating an allometric equation with credible confidence intervals for its slope and intercept. For example, taking a first example from volume allometry, if you hypothesized that special selection in humans for language ability resulted in a comparatively larger Broca’s area, it is necessary to show that the size of Broca’s area in humans exceeds its expected allometric position compared to Broca’s area in other primates (Schoenemann, 2006). A “control structure” such as primary visual cortex, a subcortical structure, or the rest of the brain cannot be used to “normalize” the volume of Broca’s area, as allometric relationships in brain volumes can be expected to be nonlinear. Broca’a area will be disproportionately large in humans versus rhesus monkeys, but it will also be disproportionately large in rhesus monkeys versus marmosets, or in horses versus sheep, where relative language competence will not apply. If Broca’s area has positive allometry compared to visual cortex, every contrast of a large and small mammalian brain will *always* show disproportionate volume increase in Broca’s area in the larger brain. Similarly, the question of whether hippocampal neurogenesis and maturation is unusually early or late in humans depends on whether the timing of hippocampal maturation deviates from its expected developmental allometry.

Inappropriate norming procedures applied to developmental timing questions will produce the identical errors to those produced by inappropriately norming allometric comparisons of volume. You cannot, for example, compare the time from birth to adolescence in chimpanzees versus humans, see that the duration is longer in humans, and conclude that human have been specially selected for a longer childhood. The duration may be entirely predictable from the time required to generate a large brain, intrinsic correlation with longevity or some other superordinate feature of life history. The “translating time” database was collected, in part, to be able to understand such comparisons in a larger cross-species context. A major surprise of this work was the extreme regularity of neural development in mammals, which in addition to the interest of the regularity alone, gives us a reliable set of brain-based benchmarks to understand the relative maturation of each species with respect to life-history events like birth or weaning (Hawkes and Finlay, 2018).

#### 1.2.2 Setting zero, or onset of neurodevelopment: birth is not a reliable indicator of brain maturation

All allometric equations have a slope and an intercept, but in developmental allometry, the intercept often suggests a real-world developmental meaning, for example, the onset of neurogenesis, or conception, or birth. Even though a real-world event like conception may appear to be a likely candidate for “zero” in an allometric equation, this must be mathematically determined, not stipulated. In “translating time”, the best fit for “day zero” to the empirically measured neuroembryological data first proved to be a point located between conception and first production of mature neurons, possibly implantation (Finlay and Darlington, 1995; Finlay et al., 2001; Workman et al., 2013). Although birth is often chosen as a natural zero in anthropological work, and especially for research on late hippocampal neurogenesis to be discussed here, for the good theoretical reason that it marks the beginning of the independent life of the organism, and for the practical reason that prenatal measurements often hard to come by, still, this choice can be very misleading when attempting to compare developmental schedules (Figure 2). We will explain the derivation of the axes and the maturational progress represented on this graph in more detail in the next section, but for the moment, the x scale, the “event scale” is a multivariate measure of overall maturational state of the nervous system, with the generation of the first neurons near “0”, with “1” corresponding to about 3 years postnatal in humans, with embryological features like achievement of 80% of adult brain volume and variable progress of myelination. The Y axis is post-conception days of development on a linear scale – on a log scale, the allometric equation of each curve plotted would become a straight line (Figure 3). Post-conception days are plotted on a linear scale in this graph to emphasize the extreme divergences in absolute days to maturity in the species plotted here.

**Figure 2.**
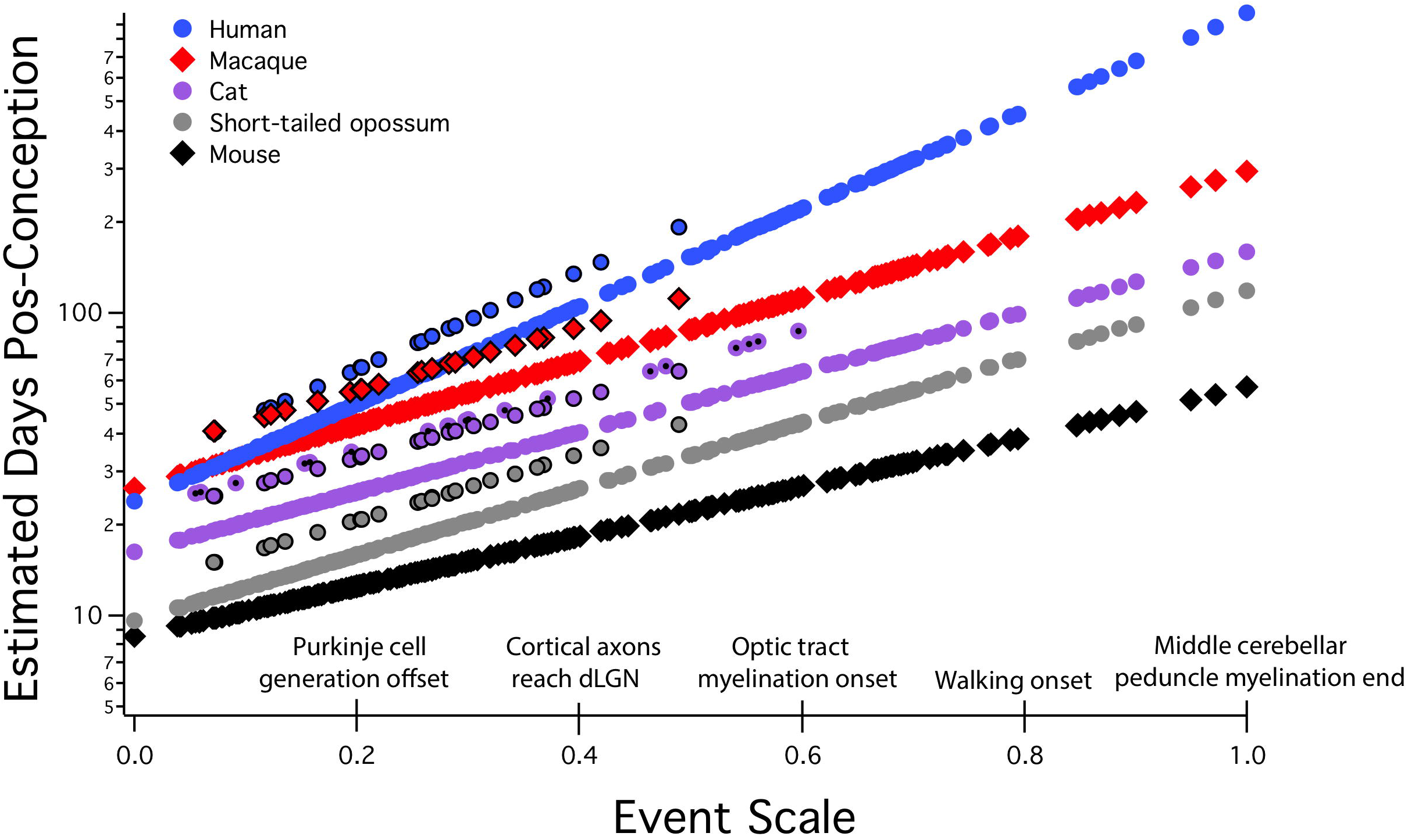
Predicted developmental schedules for human (blue circle), macaque (red diamonds), cat (purple circle), short-tailed opossum (grey circle), and mouse (black diamonds), selected from 18 species to illustrate the full range of developmental durations. This figure is modified from Workman et al., 2013. In this graph the event scale is the x-axis, to which we have added a subset of the 271 events that were observed. The event scale is a common ordering of developmental events across all species and ranges from 0 to 1. The y-axis is the estimated date of occurrence of each event in each species from conception (log scale). To determine when a particular event would be predicted to occur in any species from this graph, using the name of the event on the event scale, find where it intersects the regression line for that particular species. The y-axis value will be the predicted PC day for that event/species combination. Also represented on this graph are interaction terms for corticogenesis and retinogenesis, with interaction terms always associated with individual species. The parallel lines for a subset of events in four of the species (black bordered circles for human, macaque, cat, and possum) represent delays in cortical neurogenesis with respect to their time of occurrence in the rodent and rabbit. In the cat, a second parallel line can be seen representing the delay of retinal neurogenesis relative to the timing of other transformations (purple circle with a black dot). dLGN: dorsal lateral geniculate nucleus.

**Figure 3.**
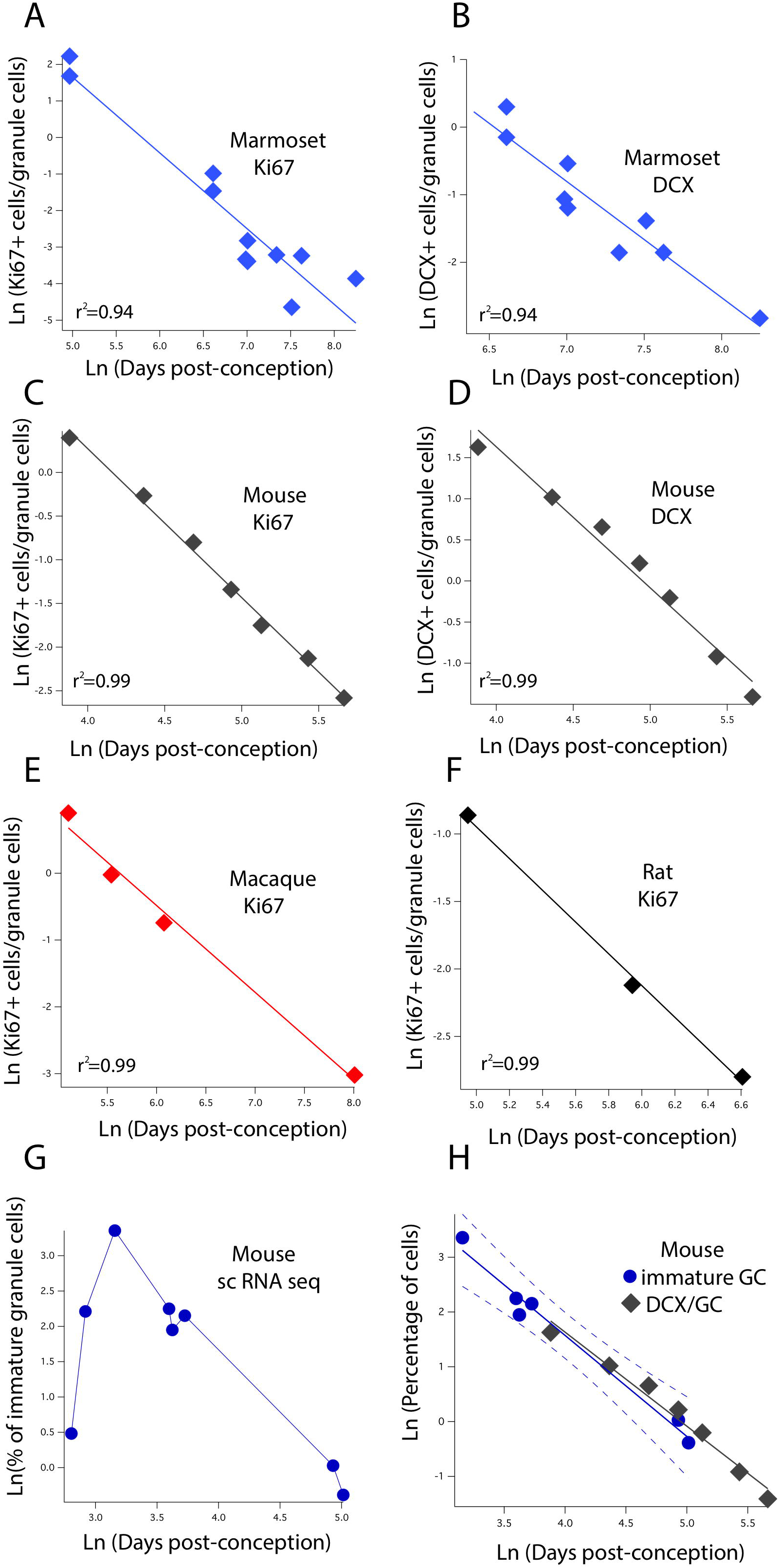
The natural-logged values of DCX+ and Ki67+ cell numbers relative to hippocampal granule cell numbers are plotted against the natural-logged values of age in days post-conception in (A-B) marmosets, (C-D) mice, (E) macaques, and (F) rats. We performed a linear regression through these values to identify when the relative number of Ki67+ cells reaches 0.7, 0.5, 0.3, 0.2 and 0.1% of the total granule cell population. We also identify when DCX cell numbers reach 3, 2.5, 2, and 1.5% of the total granule cell population. With the exception of the marmoset, the relative number of DCX+ and Ki67+ to total granule cells were averaged at each age. (G) We use single cell RNA seq to compute the number of immature granule cells relative to hippocampal neurons over the course of prenatal and postnatal development in mice. (H) Such an analysis shows that DCX+/granule cell numbers of mice fall within the 99% confidence intervals generated from single cell RNA seq data. These findings demonstrate strong concordance between methods. Data are from Merill et al., 2003; Rao et al., 2006; Jabès et al., 2010, Ben Abdallah et al., 2010; Amrein et al., 2015, and Amrein et al., 2011. Data from single cell RNA seq are from Hochgerner et al., 2018. Regressions were generated with software package IGOR.

We have stressed the importance of two basic features of developmental allometric analysis critical for interpreting the presence or absence of “postnatal” or “adult neurogenesis”. The first is obtaining developmental data from enough species to generate reliable allometric equations, and the second is locating a true “zero” from which to scale maturational events in the same equations. The Translating Time database and model can supply both necessities. Exploring “postnatal” neurogenesis in the hippocampus will be reporting on very different phenomena if mice, precocial guinea pigs, or humans are compared.

#### 1.2.3 A brief review of our specific methodology for comparing neurodevelopmental sequences across species

Over the past 20 years, a database and methodology to compare the progress of neural development across species have been elaborated (www.translatingtime.net). The multiple statistical considerations leading to this representation can be found in the series of papers detailed in the first section, and a full description of the model in Workman et al. (2013). The original purpose of this work was to describe a mammalian “Bauplan” for neural development, and thus identify deviations from this plan that might mark taxon- or species-specific alterations corresponding to evolutionary adaptations, which is exactly how we will employ it for to examine the hippocampal data we have collected. The present model includes 18 species, and 271 “events” of mixed type, including neurogenesis in particular structures or cell classes (e.g., Layer 4 of striate cortex; Purkinje cells in the cerebellum; onset of synaptogenesis in a thalamic nucleus; emergence of some minimal behavioral reactivity, and transitions capturing continuous processes such as increases in brain volume or myelination).

The model from Workman et al., 2013 is reproduced in Figure 2, and extends to a maturational stage equal to approximately 3 years postnatal in humans. Only events in brain and some early behavioral capacities are included to model the event scale and each species’ regression line – no measures of body or organ maturation or volume, or interactional, life history events like birth or weaning are included in this version. The “event scale”, which is the best order and interval relationship of the 271 distinct neurodevelopmental events in the 18 species, is fit iteratively to all the data, (x-axis, Figure 2). The speed of progress of each individual species through these events is given as a regression equation, in days on a log scale (y-axis; compare the linear scale in Figure 2 of the same functions). It is more typical to plot time on the x-axis in developmental studies, and it is important to remember this difference in representation. Days are on the Y-axis because we are interested in duration as a function of maturational state. For example, for species with different sized brains, how long will it take them to reach equivalent maturational states? The differences in each species’ slope show differences in maturational rate, with steeper slopes meaning slower progress through maturational stages in absolute time: the mouse takes only about 30 days to execute its 271 neurodevelopmental events, while the human takes 1000 days, as humans generate greater numbers of neurons and volumes of connectivity per event.

The fit of model results to empirically-measured results is astonishingly close, 0.9929, which reflects an extreme, and initially unexpected conservation of developmental sequences in mammals. Only two interaction terms are necessary to produce taxon-specific differences in these data so far, which are the black-circled points floating above the larger number of points of the corresponding color. The first term corresponds to a delay in corticogenesis in primates, some marsupial species and carnivores (n.b: this can be equally well represented as an advance in initiation and termination of neurogenesis in the “rest of the brain” --Clancy et al., 2001; Workman et al., 2013, Charvet et al., 2017ab). The second represents a delay in neurogenesis in the retina of the nocturnal cat and ferret (also owl monkey, Dyer et al., 2009). Extensions in cortical neurogenesis produce a disproportionate expansion of the cortex and, in particular, upper layer neurons in primates (Cahalane et al., 2014; Charvet et al., 2015, 2017ab), and a greater number of rods and rod-associated neurons in carnivores and owl monkeys.

##### 1.2.3.1 Birth can intersect quite different developmental events in different species

As noted earlier, birth may occur at a wide range of stages in neural development in different species. For example, cortical and cerebellar neurogenesis is ongoing at birth in some rodents, but in primates, both are largely concluded at that time. No obvious inflections, halts or accelerations near birth can be found in basic central nervous system construction. There is one event, a whole-brain surge of synaptogenesis, which appears to just antedate either birth or burrow exit in the four mammals studied to date instead of conforming the otherwise monolithic neurodevelopment program (reviewed in Finlay and Uchiyama, 2017).

##### 1.2.3.2 Other evidence for regular mammalian neurodevelopment

Empirical support for the surprising claim of an extremely conserved mammalian neurodevelopmental schedule can be found in several independent sources. Mammalian brains continue to grow after birth, and Passingham (1985) first noted that if the volume of the brain at birth is plotted against gestation length for an eclectic set of eutherian mammals, including rats, pigs and dolphins (log transformed), a straight line results, suggesting brain mass is produced generally at the same rates in all species, smaller brains simply ceasing their growth earlier (Passingham, 1985). Halley, in a much larger and more closely measured data set of changes in brain volume post conception, recently confirmed the same notion (Halley, 2016, 2017). We have also successfully modeled the development of neuron number in the cortex combining information on kinetics of neurogenesis with adult neuron numbers in multiple species (Charvet and Finlay, 2014; Cahalane et al., 2014). Other observations of single maturational phenomena give other insights, and underline further unexpected consequences of this conserved neurodevelopmental rate.

##### 1.2.3.3. Two surprising findings about precocial animals

In mammals, the onset of walking is predicted by neural maturation (which is conserved) but not birth or any known niche variable. The time of the first unsupported step is highly predictable from a developmental allometric equation derived from adult brain mass, including one interaction term slightly accelerating the time of first step for those species with a plantigrade standing position (Garwicz et al., 2009), which fits seamlessly into the translating time model. This monolithic nature of the neurodevelopmental program, and its close correlation with brain size puts an interesting constraint on precocial mammals. Relatively large-brained, precocial ungulates like sheep and elk, who must be ready to run just after birth, accomplished this evolutionarily by extending gestation and delaying birth in their large offspring to match conserved parameters of brain development. They do not selectively advance the general rate of brain maturation nor push forward the maturation of circuitry closely associated with ambulation apart from the rest of the brain, which might seem to be a less stressful solution.

A related peculiarity can be seen in precocial species with relatively small brains such as the guinea pig and spiny mouse, that are born looking and moving quite mature, furred, and with sensory systems functional. While it might seem a reasonable strategy to make the most of every possible second for brain maturation available *in utero* in precocial species, to allow fine tuning of the coordinated behavior required immediately after birth, the conserved pace of brain maturation seems to rule this out. Since these animals must also produce large, mature bodies, which appear to require more time than the brain, the onset of neural development as marked by the first postmitotic neurons is substantially *delayed,* not stretched to fill the available time, allowing somatic maturation a head start (Workman et al., 2013). We will discuss whether a similar situation is present in marmosets, born with some precocial features.

### 1.3 Applying “Translating Time” to the question of late hippocampal neurogenesis

The first reports of neurogenesis in adult humans and other mammals produced much excitement, in that it contradicted the central dogma that no new neurons, are generated in adulthood and offered a possible avenue for brain rehabilitation and repair. At first, the presence of new neurons was reported widely throughout the forebrain, but in time, unambiguous neurogenesis was finally limited to two locations, the hippocampus and the olfactory bulb via the “rostral migratory stream”, mostly from work in rodents, but with confirmation in humans (Ming and Song, 2005). Recently, however, the existence of significant adult hippocampal neurogenesis has been questioned (Dennis et al., 2016; Kempermann et al., 2018; Andreae et al., 2018; Sorrells et al., 2018; Lee and Thuret, 2018). A report by Sorrells et al. (2018) concluded that neurogenesis in the human hippocampal dentate gyrus drops to undetectable levels during childhood, suggesting that human hippocampal neurogenesis is unlike that of other mammals (Knoth et al., 2010). A concurrent study (Boldrini et al., 2018) also investigated adult neurogenesis in the human hippocampus (14 to 79 years of age) and contradicts the first study. Using quite similar methodologies, the second group argued that adult hippocampal neurogenesis is in fact present throughout the lifespan. In such cases of contradiction, consultation with the animal model literature is of major help. A problem that has plagued this work is the absence of a robust and reliable way to compare time courses of events in different species. Adult hippocampal neurogenesis of any species could represent the tail end of a normal embryonic period of neurogenesis, or a truly indeterminate phenomenon, as is seen in virtually all non-mammalian vertebrates, or perhaps a targeted rekindling of neurogenesis for a particular purpose in adulthood. Because of the methodological similarity of the two studies, it may not be possible to rule in favor of one or the other on reported evidence, but a better idea of where errors might lie is a natural outcome of quantitative developmental modeling.

#### 1.3.1 Specific objectives 1: extending the translating time model and representing species on a common scale

As we described previously, mammalian species vary in the length of both neural and somatic development, the positioning of birth with respect to neural maturation, and the relative length of neurogenesis in different structures. Comparing humans to macaques and mice, human neurodevelopment is much longer (duration correlating close with brain volume, as does the duration of lifespan). Humans are born at a slightly earlier stage of neural maturation than macaques, and at much later stage than rats and mice. Rhesus monkeys and humans also curtail neurogenesis in limbic structures relatively earlier than rodents (Workman et al., 2013), corresponding to the fact that as limbic structures are systematically relatively smaller (that is, scale with a smaller exponent) in primates compared to rodents (Reep et al., 2007). The translating time model at present does not have good data representation for late developmental stages to allow close comparisons in adulthood. We are therefore adding new data, and one new species to extend the model farther into the lifespan, but without any substantive change in its basic structure. We find appropriately-transformed envelopes of neurogenesis across species to be very similar, and continuous.

#### 1.3.2 Specific objectives 1: Closer examination of human hippocampal neurogenesis and the problems of detecting non-scaling cellular events in a nonlinearly scaling lifespan

We consider the allometric nature of developmental schedules in humans to identify how hippocampal neurogenesis should vary if the duration of hippocampal neurogenesis in humans is similar to that of rodents. Further, the ability to align timetables allows us to investigate an intrinsic problem of detection of a cellular signal in scaling situations, which is that organismal variables of size and duration show robust scaling, but cellular phenomena like action potentials, the length of the cell cycle and so forth rarely do. A rat may expect to live around 700 days post-adolescence, while an approximate comparable human figure is 25,000 days. If the cellular processes associated with an occasion of neurogenesis are transitory, and almost certainly do not scale with lifespan, the probability of simple detection falls radically in the long lifespan. We will discuss this aspect of scaling both as a methodological problem, and as a question about the importance of extremely low-probability events.

## 2.0 Materials and Methods

### 2.1 Species and sources

In order to extend the current neurodevelopmental model to later developmental stages, we added some additional data on the timing of developmental milestones in two rodent species (i.e., rats, mice), and 3 primate species (i.e., macaques, marmosets, humans). “Developmental events” capture rapid transformations, such as onset of neurogenesis or any other process, or arbitrary divisions of continuous processes into epochs (e.g, 20%, 40%, 60% and 80% of a structure’s adult volume). Examples of developmental events include birth-dating of cell types, synaptogenesis, myelination, changes in protein and RNA expression. The new types of data added were those capturing temporal changes in cell proliferation from markers (i.e, DCX, Ki67) in the hippocampus. We only include developmental events present in at least two species, and at least one rodent species.

We identified variation in proliferative and newly born neuron numbers over the course of prenatal and postnatal development in primates and in rodents. More specifically, we collected previously published data where the number of Ki 67+ (proliferative) and newly born (DCX+) cells relative to the total number of hippocampal granule cells was quantified at several stages of development in rodents and in primates. We defined as epochs when Ki67+ cells decline to 2%, 0.7%, 0.5%, 0.3%, 0.2%, and 0.1% of total granule cells in primates and rodents. We also identified when the number of DCX+ cells reach 3, 2.5, 2, 1.5, 1, and 0.5% of total granule cells in rodents and in primates. To do so, we fit a linear regression between the natural-logged values of age and the relative number of cell markers to compare the duration of the decline in late hippocampal neurogenesis between primates and rodents (Figure 3). We only selected age ranges in which there is a sharp decline in the relative number of Ki67 and DCX+ cells over time as assessed on a natural-log scale. This permits fitting a linear regression through the data for each species (Figure 3). These data are from Merill et al., 2003; Rao et al., 2006; Jabès et al., 2010, Ben Abdallah et al., 2010; Amrein et al., 2015, Amrein et al., 2011, and Hochgerner et al., 2018. For rats, we considered the number of Ki67+ and DCX+ cells from Rao et al., 2006 and total granule cell numbers from Merill et al., 2003. We consider studies that normalize the total number of proliferative and immature cells relative to total granule cells rather than those that consider the number of proliferative and immature neurons per mm^2^ of tissue.

We consider developmental transitions as the emergence of “plateaus” in the expression of multiple genes in single structures. We identified such plateaus in RNA expression from RNA sequencing data of bulk from the hippocampus in both species (Iacono et al., 2017). We identified when expressed genes reach a plateau in their expression across 14,417 orthologous genes as defined by the mouse genome database (Smith et al., 2018). We used a non-linear model with the software package R (easynls, model 3). Only orthologous expressed genes were considered. Age ranges were constrained to vary between 101 to 999 days in humans (n=10) and between P1 to P30 in mice (n=15) to compare roughly equivalent developmental time windows across these two species. We used normalized RNA sequencing expression made available by the Allen brain Institute. RNA expression from mice hippocampi was obtained from Iacono et al. (2017; GEO: GSE79380). We selected only those models with p values of coefficients less than 0.05 in humans and mice. This resulted in 34 genes in which plateaus were identified in both species. We averaged the age in which plateaus in RNA expression we identified in both species and include these data as one developmental event.

### 2.2 Developmental timing in marmosets

We gathered available data on the timing of early neurodevelopmental events for the marmoset as we had done for other species. We matched our previously collected database on developmental event definitions, principally using anatomical changes from structural MRI scans (Hikishima et al., 2013), spatiotemporal changes in gene expression, as well as anatomical transformations from the literature. Examples of developmental events include morphological events such as first observation of retinal axons in the optic stalk, or when neurofilament heavy polypeptide (NEFH) expression emerges in the cortex. Because the marmoset is increasingly used as a model organism, we expect this inclusion to be useful past this study alone. To compute the timing of developmental events from MRIs, we noted the earliest age in which a event had occurred and the latest age in which the event had not yet occurred, as we had done previously (Charvet et al., 2010; Workman et al., 2013). In total, we include 29 events for marmosets.

### 2.3 Statistical analyses

We include 213 developmental events from Workman et al., 2013 and Charvet et al., 2017b, eliminating events capturing the timing of cortical neurogenesis because cortical neurogenesis is extended in primates compared with rodents. We only included developmental events present in at least two of the species. Of the 213 events, 47 represented events or stages in limbic system development, including neurogenesis timing as well as the emergence of axonal pathways of limbic structures. We added 22 developmental events, 14 of which that capture the decline in late hippocampal neurogenesis (6 Ki67, 8 DCx; Figure 3).

A new 0-1 “event scale” was fit linearly to span this extended range, by subtracting the timing of each developmental event from the earliest event and divided these values by the difference between the latest event and the earliest event. We fit a linear regression through log-transformed values for each species against the event scale. We use the fitted values from the regression of human developmental event timing versus the event scale to predict the timing of late stages of human hippocampal neurogenesis timing. We omitted developmental events capturing cortical neurogenesis because a subset of cortical cell types are generated later than expected in primates (Clancy et al., 2001; Charvet et al., 2017ab), which may increase error when predicting the duration of hippocampal neurogenesis across species.

We tested whether hippocampal neurogenesis occurs earlier than expected given the timing of other developmental events. We fit a linear model with the event scale as a continuous variable and developmental event timing as the response variable. To test whether limbic structures undergo neurogenesis earlier than expected relative to the timing of other events, we classified neurogenetic events as limbic or non-limbic. We tested whether the “limbic factor” as well as the interaction between the event scale and the “limbic factor” would account for a significant percentage of the variance.

### 2.4 Single cell RNA sequencing to identify adult hippocampal neurogenesis

Because adult neurogenesis has recently been disputed in humans (Sorrells et al., 2018), we investigated whether adult neurogenesis could be observed from single cell RNA sequencing data extracted from the human hippocampus and prefrontal cortex aged 40 to 65 (Habib et al., 2017). We computed the relative number of cells expressing neural progenitor markers (DCX+, SOX2+, DPYSL3+) relative to the number of cells expressing PROX1+ in humans. We select PROX1 as a marker for granule cells because it is expressed by hippocampal granule cells but not by other cell types in the cortex. That is, the expression of PROX1 from bulk samples is higher in the hippocampus than in other cortical regions (Figure S1A) and PROX1 is expressed by hippocampal granule cells but not by isocortical cells (Figure S1B). We selected SOX2, DCX, and DPYSL3 (aka TUC-4) because they are markers of immature neurons (Ngwenya et al., 2006; Cipriani et al., 2018). We considered PROX1 to be expressed if the gene count was greater than 0. To identify whether DCX+, SOX2+, and DPYSL3+ collocate with PROX1+ cells above chance level, we randomly reassigned PROX1 expression to different neuronal types 1,000 times. We then computed the number of DCX+, SOX2+, and DPYSL3+ cells relative to the number of PROX1+ cells. We assess whether the relative number of DCX+, SOX2+, +, and DPYSL3+ falls above the 95% confidence intervals generated from 1,000 permutations. Such an analysis permits investigating whether the number of immature neurons is present above chance level. Because we are focused on whether new neurons are generated in the adult hippocampus, we do not include cells belonging to clusters previously identified as glial, astrocytic, microglia, and endothelial. Data are from DroNc-Seq generated by Habib et al., 2017.

## 3.0 Results

### 3.1 Initial characterization of late hippocampal neurogenesis

The initial step is to characterize how the number of dividing progenitors (Ki67+ cells) and immature neurons (DCX+) relative to granule cell numbers vary with post-conceptional day. Figure 3 (A, C, E, and F) show the measured values of Ki67+ cells expressed as a percent of total granule cells versus days post-conception for the marmoset, mouse, macaque, and rat. Frames B and D show DCX+ labeled granule cells for marmoset and mouse only, again as a percent of total granule cells. Both scales are natural log scales, and the durations spanned vary considerably, from approximately 50 to 250 days post conception in the mouse, versus approximately 150 to 3,000 days postnatal in macaque and marmoset. This enables calculating when the percentage of Ki67+ to total granule cells reach 2%, 0.7%, 0.5%, 0.3%, 0.2%, and 0.1%, and when the percentage of DCX+ to total granule cells reach 3%, 2.5%, 2%, 1.5%, 1%, and 0.5% in each species. The range for each species was constrained so that the natural-logged values of the relative number of Ki67+ and DCX+ to total granule cells systematically decline with age. This approach permitted fitting a linear regression through the data.

### 3.2 Addition of declining hippocampal neurogenesis values into the overall maturational event scale

In Figure 3, hippocampal neurogenesis indicators are described with relation to post-conception day in each species, but we would like to know how the decline in hippocampal neurogenesis relates to the common progress of brain maturation across species. Two ways of presenting “translating time” data can be used. In Figure 4A, the new data on late hippocampal neurogenesis for marmoset, macaque and mouse, and the single rat point are plotted against the common “event scale”. This type of data representation is optimal for visualizing overall slope and intercept similarities and differences between multiple species. As expected, the species with long early neurodevelopment periods show longer periods of adult hippocampal neurogenesis. No truncations, breaks or sudden accelerations in any particular species are in evidence, though there are interesting differences in the maturational path in marmosets versus macaques we will address subsequently.

**Figure 4.**
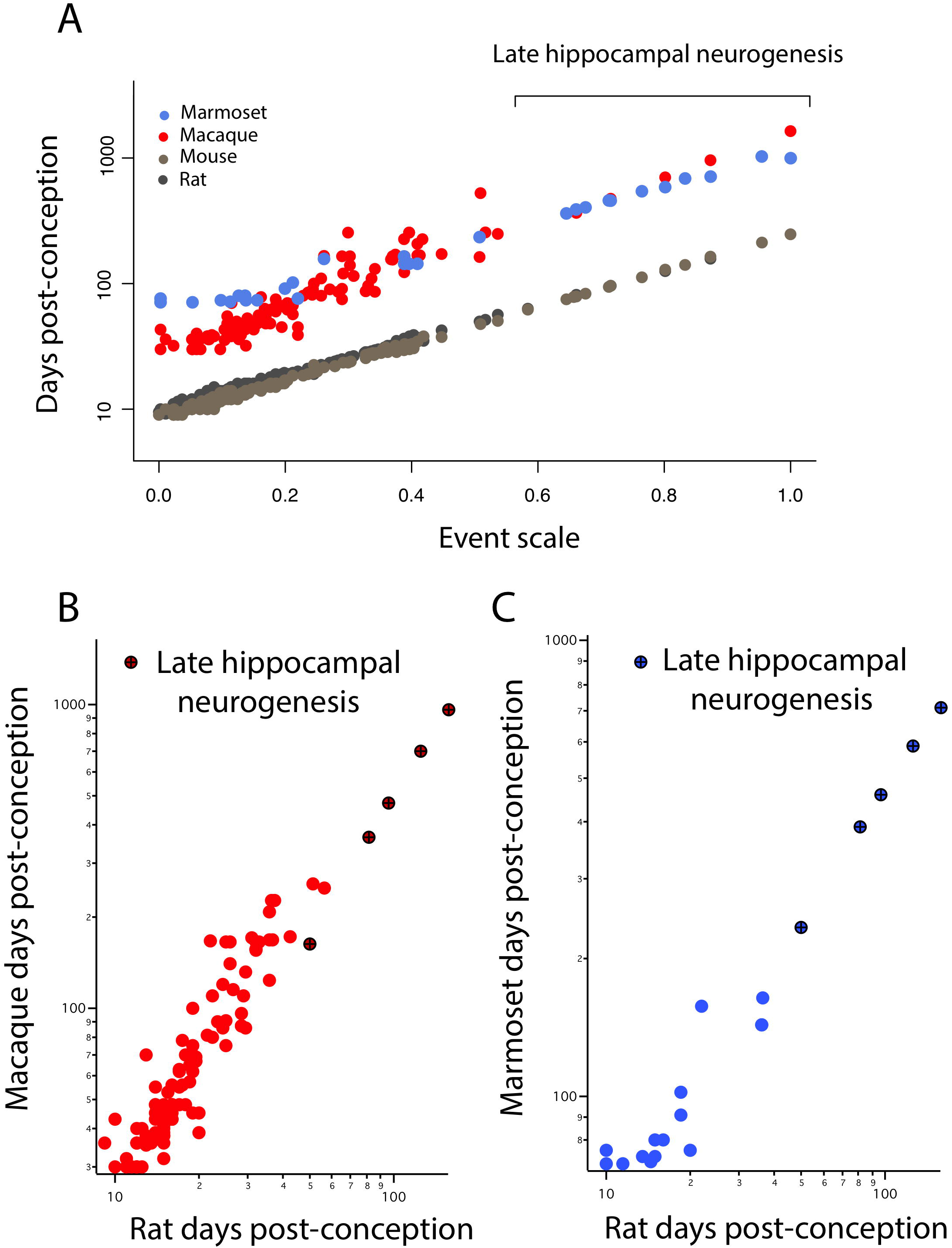
(A) The timing of developmental events is plotted against an event scale in macaques, marmosets, rats, and mice. The timing of hippocampal neurogenesis was extrapolated from regressions capturing the decline in the number of DCX and Ki67+ cells relative to hippocampal granule cell numbers in primates and rodents. The timing of developmental events in macaques (B) and marmosets (C) are plotted against those in rats. (B-C) Developmental time points capturing late stages of hippocampal neurogenesis are highlighted (x).

It is also possible to use the translating time scale to express the events of one species in the time frame of a second species, “translate” the approximately 130 modeled days of the macaque to the 50 days of a mouse, which facilitates close comparisons of delay or advance of any class of events between the selected species (Figure 4, B and C). For example, comparing nocturnal to diurnal mammals, the rods and other rod-related cells of the retina are generated later in nocturnal mammals, which would be visible in graphs like 4B and C as an elevation of rod-related points (nocturnal animals on the Y axis) (Dyer et al. 2009; Workman et al. 2013). In this case, we look for a difference in the implied intercept or slope of the “late hippocampal neurogenesis” points to determine if they show any signs of systematic variation from the common developmental scaling function. No such differences are apparent.

#### 3.2.1. Marmoset developmental timing

Early in development, equivalent events in marmosets occur later than in macaques (Figure 4A). At later time points, equivalent events occur earlier in marmosets than in macaques. A linear model with the event scale as a continuous variable and the logged values of developmental event timing as the predictor shows that the slope is lower in marmosets (y=1.27x+1.73, slope SE=0.037, intercept SE=0.02, R^2^=0.976, p<2.2e-16) than it is in macaques (y=1.89x+1.44, slope SE=0.056, intercept SE=0.016, R^2^=0.903; p<2.2e-16). In other words, marmosets initiate neural development late with respect to conception, close to day 90 compared to day 35 in macaques, but then progress through developmental events faster than macaques, producing a smaller brain by the end of neural development. The consequence of this late, accelerated developmental trajectory is that hippocampal neurogenesis wanes earlier in marmosets than in macaques. This is similar to the pattern previously observed in precocial mammals like guinea pigs, spiny mouse and sheep (Workman et al., 2013) where neural development is delayed with respect to conception later, but once initiated, proceeds at a faster rate than in a number of altricial species.

#### 3.2.2. Somewhat earlier termination of limbic neurogenesis in the macaque

The large sample size in macaques allows us to test whether limbic neurogenesis occurs earlier relative to the timing of other events in macaques (Figure 5). To that end, we fit a linear model with the logged values of developmental event timing as the predictor, the event scale as a continuous variable and a discrete categorical variable that classifies neurogenetic events as limbic or not. We also tested whether the interaction between the “limbic” factor and the event scale accounts for a significant percentage of the variance. The fitted model accounts for a significant percentage of the variance in developmental event timing for macaques (F=416.7; R^2^=0.91). The limbic factor is not significant (F=2.065; p=0.15) but the interaction between the limbic factor and the event scale is significant for macaques (F=11.92; p<0.05). These data demonstrate that the slope of the natural-logged values of late hippocampal neurogenesis versus the event scale is lower than expected in macaques considering the timing of other developmental events. In other words, hippocampal neurogenesis may cease slightly earlier than expected in macaques compared with rodents.

**Figure 5.**
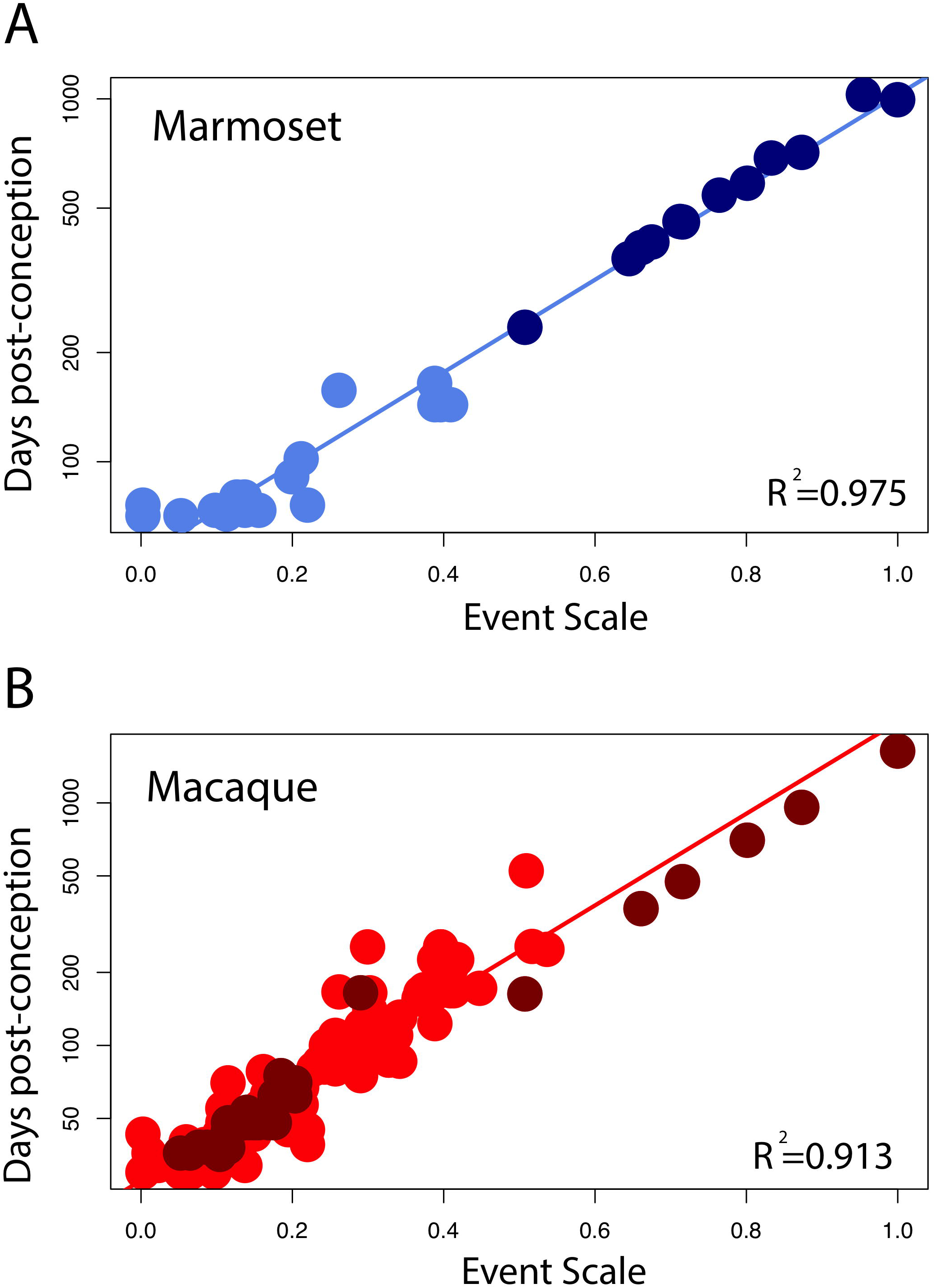
Developmental milestones are plotted against the event scale in marmosets (A) and macaques (B). Milestones that capture the timing of limbic neurogenesis are in dark blue (marmosets) and in dark red (macaque). We fit a linear regression through the logged values of the timing of developmental milestones against the event scale. Late hippocampal neurogenesis consistently falls below the regression. In other words, late hippocampal neurogenesis may occur earlier than expected given the timing of most developmental milestones in macaques.

#### 3.2.3 Hippocampal neurogenesis in humans

For humans, a linear regression of the timing of reported developmental milestones versus the event scale computed for humans by the translating time model (Workman et al., 2013) is plotted for the reduced dataset we used in this model, as a visual check and demonstration of the predictability of human data points, in Figure 6A. No new data are introduced in Figure 6A; its intention is only to show the baseline variability against which we might introduce and compare other data. (y=2.44x+1.53; slope SE=0.12; intercept SE=0.04, R^2^=0.85, p <2.2e-16). We then extrapolated predicted values from the linear model to see how late stages of hippocampal neurogenesis should vary if the timing of hippocampal neurogenesis were conserved across humans and mice (Solid lines, Figures 6B and 6C) According to these predictions from mice, human hippocampal neurogenesis as assessed from the relative number of Ki67+ and DCX+ cells should drop sharply between prenatal stages up until to 8-26 years of age and subsequently remain relatively invariant at later time points (Figure 6B). More specifically, the percentage of Ki67+ to total granule cells should drop up until about 8 to 26 years of age (post-conception day 3,000 to 10,000; Figure 6) and remain relatively invariant thereafter. Similarly, the relative number of DCX to total granule cells should drop from birth to about 8 to 26 years of age (post-conception day 3,000 to 10,000; Figure 6C.

**Figure 6.**
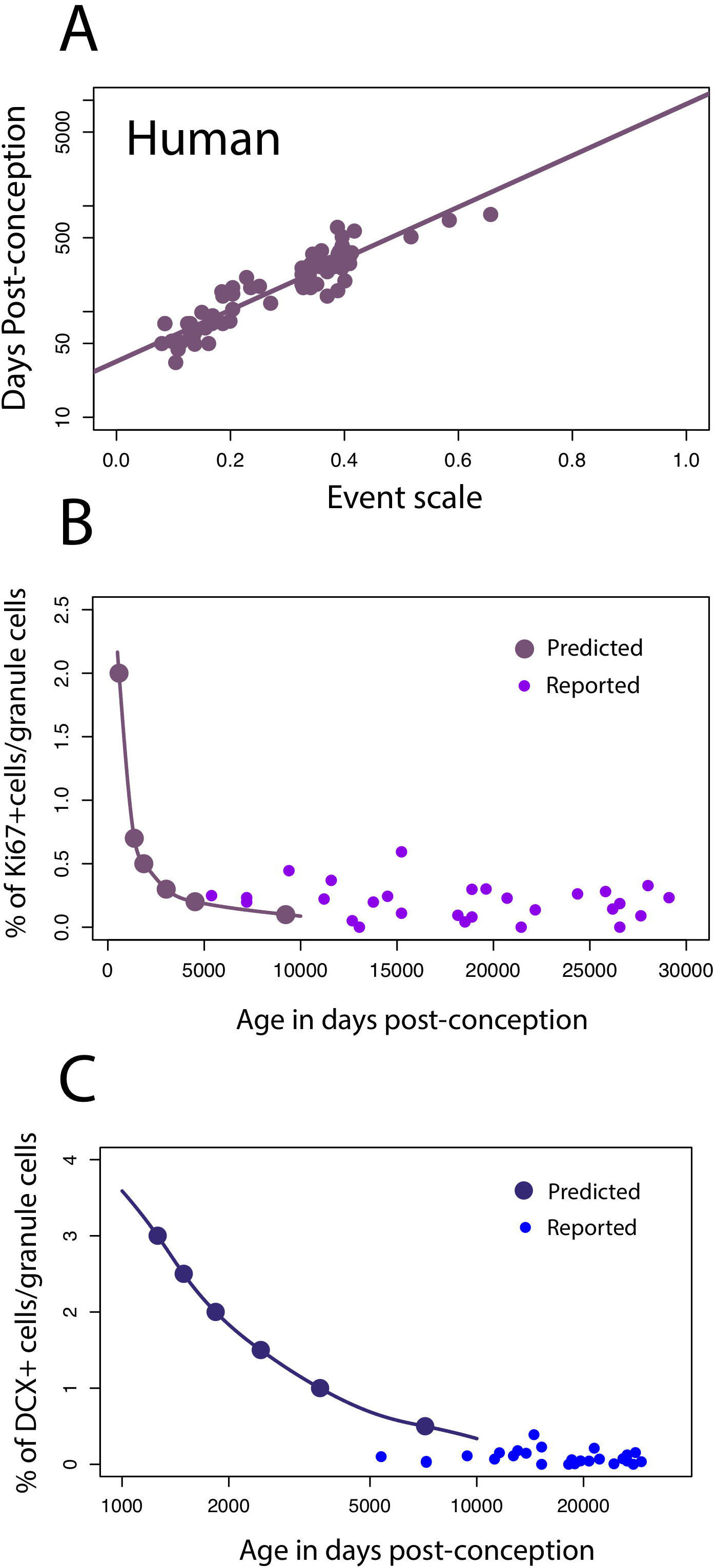
(A) The timing of developmental milestones in humans are plotted against the event scale. We fit a regression through the timing of developmental transformations against age in days post-conception. We use this regression to predict the decline in late hippocampal neurogenesis in humans. (B_C) The number of Ki67+ (B) and DCX+ (C) cells relative to the total granule cell population is predicted to decline sharply during childhood in humans.

On these predicted functions, we overlay the number of DCX+ and Ki67+ cells compared to total granule cells as reported by Boldrini et al., 2018. Our predictions are very generally consistent with those of Boldrini et al., in that we predict that both markers should remain relatively invariant from 8-26 years of age onward, but the added data do appear more variable, and the Ki67+ cell numbers higher than would be expected. Other data potentially addressing this timetable, that of Sorrells et al. (2018), could not be plotted against this representation because number of proliferative cells/mm2 was assessed rather than relative to the total granule cell numbers determined in the rodent studies.

To investigate whether hippocampal neurogenesis timing in humans should deviate from that of rodents, we compare temporal changes in DCX expression in humans and mice. These data offer a slightly different perspective on the temporal pattern of late stages of hippocampal neurogenesis between species. We first note similarities between DCX RNA expression and the relative number of immature hippocampal granule cells assayed from single cell RNA sequencing data (Figure S2). A qualitative investigation of DCX expression from multiple datasets in mice suggests that RNA sequencing from bulk data mirrors the temporal changes in the relative number of immature granule cells. As the relative number of granule cells declines in mice, DCX expression from bulk samples also declines sharply. At roughly 38 to 50 days post-conception, the relative number of immature granule cells varies relatively compared to earlier time points. That is, DCX expression is relatively invariant from 1 month to 4 months of age in mice. In humans, DCX expression also decreases from prenatal time points up until around post-conception 316 (50 days after birth) and subsequently remains relatively invariant. According to the translating time model, 38 to 50 days post-conception in mice is roughly equivalent to 445 to 700 days post-conception in humans. In other words, the end in the abrupt decline in DCX expression might occur slightly earlier than expected in humans.

Because the presence of hippocampal neurogenesis has recently been questioned, we investigate whether hippocampal neurogenesis can be observed from single cell RNA sequencing obtained from the human hippocampus and prefrontal cortex (Figure 7A). We compare the number of cells expressing DPYSL3, DCX, and SOX2 relative to the number of PROX1 cells (Figure 7 B-E). PROX1 is used as a marker of hippocampal granule cells and its expression is observed in previously identified excitatory hippocampal granule cells (cluster 8) and GABAergic cells (cluster 7). We computed the number of DCX+, SOX2+, and DPYSL3+ cells relative to the number of PROX1+ cells. We assess whether these values lie above chance level by comparing these values to those generated by permutation-based significance thresholds. Such an analysis shows that the SOX2+ and DPYSL3+ cell numbers relative to PROX1+ cell numbers occurs above the 99% confidence intervals of distributions generated from permutations (Figure 7 F-H). However, the number of DCX+ to PROX1+ cells falls within the 99% confidence intervals generated from permutations. These findings suggest that human hippocampal neurogenesis is present at low but detectable numbers in the adult human brain but that DCX expression may drop to such low levels in adulthood that human hippocampal neurogenesis may be difficult to conclusively identify with DCX RNA expression.

**Figure 7.**
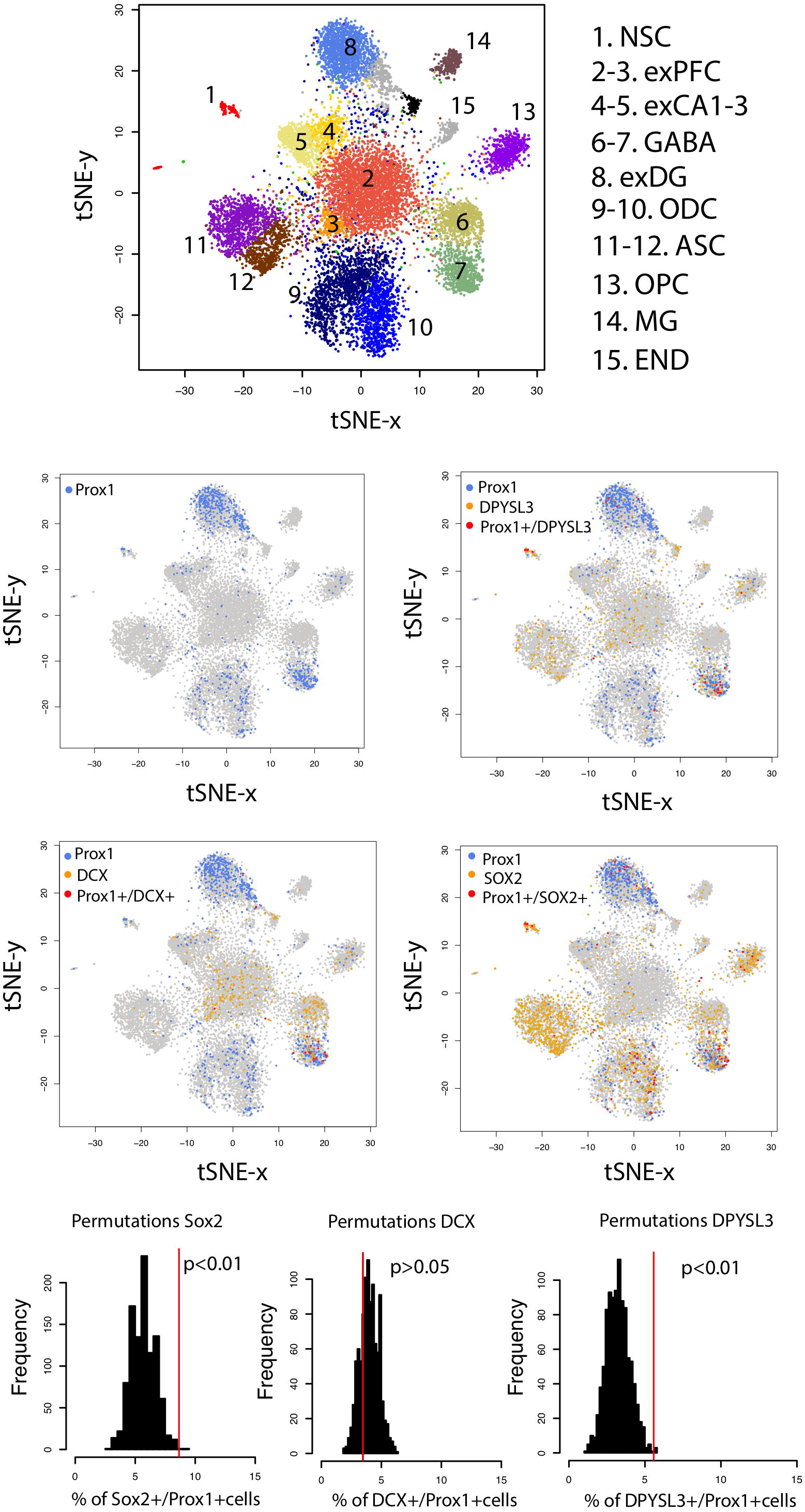
(A) t_SNE plots of RNA expression from single cells extracted from the hippocampus and the prefrontal cortex of the human brain identifies clusters of cell populations. (B) PROX1 is used as a marker of hippocampal granule cells and its expression is observed in previously identified excitatory hippocampal granule cells (cluster 8) and GABAergic cells (cluster 7). (C) PROX1 co-localizes with genes expressed by immature neurons ((D) DPYSL3 (E) DCX (F) SOX2), which suggests that new granule cells are born in the human hippocampus. To identify whether DCX+, SOX2+, and DPYSL3+ collocate with PROX1+ cells above chance level, we randomly reassigned PROX1 expression to different neuronal types 1,000 times. We then extracted the number of DCX+, SOX2+, and DPYSL3+ cells relative to the number of PROX1+ cells 1000 times (F-H). Such an analysis shows that the SOX2+ (F) and DPYSL3+ (H) cell numbers relative to PROX1+ cell numbers occurs above the 99% confidence intervals generated from permutations. However, the number of DCX+ to PROX1+ cells falls within the 99% confidence intervals generated from permutations (G). We removed cells belonging to clusters 9-15 from the analyses because they are identified as glia, astrocytic, microglia, and endothelial cell populations. We omit these cell types because our analysis is focused on identifying whether neurons rather than glial and endothelial cells are generated in the adult hippocampus. Data and identified clusters are from DroNc-Seq generated by Habib et al., 2017. Abbreviations: NSC: neural stem cells; exPFC: excitatory neurons in the prefrontal cortex; exCA1-3: excitatory neurons in CA1-3; exDG: excitatory neurons in dentate gyrus; ODC: oligodendrocytes; ASC: astrocytes; OPC: oligodendrocyte progenitors; MG: microglia; END: endothelial cells.

## 4.0 Discussion

### 4.1 Late hippocampal neurogenesis as an extension of development

When the dates and magnitudes of the long tail of declining late hippocampal neurogenesis are represented on the common maturational scale of the translating time procedure, it is clear that these events are continuous with early hippocampal neurogenesis, with little or no convincing evidence or hints of breaks or inflections. The translation of a maturational state to a particular duration of development is consistent with the normal translation seen in smaller versus larger brains.

A structural correlate of duration of neurogenesis in the embryonic brain lends additional support to the conclusion that late hippocampal neurogenesis is an aspect of developmental neurogenesis in the brain. The embryonic brain first appears as a plate, with its caudal-to-rostral dimension comprised of repeating segments, the familiar spinal segments which undergo relatively little reorganization from embryo to adult, rhombomeres to the level of the midbrain (Lumsden, 1996), and prosomeres in the telencephalon (Puelles et al., 2013; Albuixech-Crespo et al., 2018). The rhombomeric and prosomeric segments have repeating structural similarities, but undergo prolonged neurogenesis compared to the spinal cord, producing major changes in their appearance due to simple mass of neurons and neuronal migration. Important for the present purposes, the basal-to-alar dimension of the original neural plate is also a gradient of duration of neurogenesis, shortest medially and longest laterally corresponding to, but not completely accounting for the number of neurons in each segment derived from each germinal position (Finlay et al., 1998; Workman et al., 2013) At the most lateral margin of the most anterior segments that produce the pallium, we find the zones that generate the olfactory bulb, the hippocampus, and the neocortex. This region collectively generates neurons for the longest duration, the first two continuing to add neurons well past the early developmental period. Thus, extended neurogenesis is a feature of the embryonic origin of the hippocampus, not a feature applied to an unpredictable location.

#### 4.1.1 Limitations of the database

Several caveats are in order, some about the translating time approach in general, and some about the particular procedures we used to incorporate this atypical corpus of data. While the translating time database is presently the only source integrating multiple aspects of developmental information over a large number of species, from a phylogenetic perspective, those species are anything but systematically or randomly chosen, featuring a large number of rodents and marsupials, relatively few primates, with the first New World primate appearing with this article, few large ungulates or carnivores and no cetaceans. As additions of new taxonomic groups or functionally defined groups, such as the precocial mammals in Workman et al. (2013) typically reveal new ways that neural development can be altered, blanket statements about “mammalian” neurogenesis should be avoided. So far, however, a few generalities can be made. Neurogenesis can begin very rapidly after the completion of the early germinal tissue, or it can be delayed while other tissues begin proliferation as is seen in some precocial mammals, but it always moves *en bloc*, and we have never observed breaks introduced into the overall sequence, as none were observed in this analysis. The onset and offset of neurogenesis in identified groups can be shifted, most frequently seen for the limbic versus neocortex shift described earlier, a neural variation extending back to sharks and rays (Finlay and Darlington, 1995; Reep et al., 2007, Yopak et al., 2010). Finally, while duration of neurogenesis is a very important aspect of brain evolution, it is important to keep in mind it is not the only source of variation, with medial-lateral axis location, for example, only accounting for about 50% of the variance in neuron number (Finlay et al., 1998).

We estimated the relative timing of the decline in hippocampal neurogenesis not by a complete recomputation of the model to include the new observations, but rather by extrapolating the former model, duration extending almost by a factor of 2 in mice and more in the larger species, a substantial amount. It is possible that this procedure could mis-estimate the slope of the decline fairly substantially, but it seemed reasonable to attempt a first description. We did note that macaques appeared to begin initial hippocampal neurogenesis slightly earlier and end earlier than expected given the timing of surrounding, non-hippocampal events. Ideally, other late developmental events should be used to anchor these observations, at which point the overall model will be recalculated, but defined points become harder to identify in later development. Continued reduction of neuron density in most structures in later development as well as spatiotemporal changes in RNA expression are potential candidates, but these approaches have rarely been employed systematically across a broad range of species.

### 4.2 Developmental timing in marmosets

The inclusion of marmosets in the present study was intended to allow better comparisons between primate species, particularly because information on late hippocampal neurogenesis was available for it. We were somewhat thwarted in this enterprise, however, because we did not observe the simple translation for production of a smaller brain expected from the pattern laid out in rhesus macaques. Rather, early developmental events were delayed with respect to conception, then maturation proceeded rapidly, consistent with the marmoset’s smaller brain, and finally, late developmental events occurred earlier than predicted. A delay followed by rapid maturation was a pattern we had observed before, however, in precocial rodents and ungulates (Workman et al., 2013). Why the rate of neural development does not simply slow to take advantage of the extra time *in utero* is unclear. We have not yet observed any case of slowing of rate of neural production in eutherian mammals, although marsupials generate neural tissue at a slower rate overall (Darlington et al., 1999). Marmosets do have small brains compared to macaques, and perhaps to have time to generate the body, it is necessary to delay the onset of generation of the brain, to avoid producing a post-mature brain while still *in utero* if no change in its rate of development is possible. In a prior study of retinal neurogenesis in the owl monkey, *Aotus*, compared to the capuchin monkey *Cebus apella*, we had notice that gestational lengths were longer than we had anticipated from earlier work in Old World monkeys (Dyer et al., 2007). The reason for this potential difference in life history parameters will require more observations in the marmoset, and other New World monkeys as well.

#### 4.2.2 The timing of late stages of human hippocampal neurogenesis and the problems of detecting rare events

We used the timing of developmental transformation across non-human mammalian species to predict the timing of late stages of hippocampal neurogenesis in humans. If the timing of hippocampal neurogenesis is conserved across humans, marmosets, and rodents, hippocampal neurogenesis as assessed from the relative number of DCX+ and Ki67+ to total granule cells should drop sharply up until 8-26 years of age and remain low and invariant at later time points. Sorrells et al., 2018 showed that the relative number of Ki67+ and DCX+ cells drop sharply during childhood up until 7 to 13 years of age, consistent with our predictions in humans. The question is whether human hippocampal neurogenesis ends earlier than expected given the developmental allometries of late developmental events. It is presently difficult to determine whether human hippocampal neurogenesis does indeed deviate from predictions generated from rodents. However, the work of Boldrini et al., 2018, using similar techniques demonstrates low and invariant human hippocampal neurogenesis between the ages of 14 to 79 years, but a markedly higher incidence than the prior studies.

We found inconsistent evidence for late hippocampal neurogenesis in humans within our own study. Our analysis of RNA sequences from single cells showed that the relative number of immature neurons to PROX1 (i.e., a marker of granule cells) were observed at greater than chance levels in adult humans. However, the relative number of DCX+/PROX1 is unusually low and fell below chance levels as assessed from our permutation-based significance thresholds. Whether DCX expression is expressed at high enough levels for it to be reliably detected in the adult human brain is unclear. Although the number of potential confounds to detection of immature neurons are many, including retention of immature neuron morphology in displaced populations until adulthood (Piumatti et al., 2018); unusual levels of genetic variation (Kaushal et al., 2003) as well as all of the problems of processing of human tissue, developmental scaling plays a role as well.

As mentioned at the outset of this discussion, different components of the same tissue may scale in altogether different ways with respect to brain size and developmental duration. As a rule of thumb, with the usual number of exceptions, cell-based properties do not scale with brain size or developmental duration. As cells essentially depend on diffusion for many critical metabolic factors, in at least one plane of section, neuron diameter cannot scale with brain size (long axons, but not fat ones, are acceptable). Oxidative metabolism, action potentials and most other cellular processes ignore animal mass. How about the cell cycle? The cell cycles of initial neurogenesis take similar amounts of time in small and large brains (Takahashi et al., 1994; Charvet and Striedter, 2008). The duration of the cell cycle becomes longer and longer as maturation proceeds, not by uniform elongation of every part, but particularly the quiescent period; in addition, fewer and fewer cells contribute to the cell cycle (Takahashi et al., 1994; Kornack and Rakic., 2008; Charvet and Striedter, 2008). We have claimed, though, that the termination of hippocampal neurogenesis looks quite similar in its overall envelope across the rodents, monkeys and the human we measured. This is with respect to these animals’ maturational state, however, not their age in days. A back-of-napkin calculation of the duration of equivalent maturational periods from early “childhood” to death would be about 700 days for rats and about 25,000 days for a human. The initial spatial densities for Ki67 and DCX+ are roughly similar (Figure 3). It seems unlikely that generating a neuron would require 36 times longer in humans, or that the features of a young neuron would persist a similar long duration. On the other hand, the migration and integration of new neurons into circuits has been reported to take very much longer in adults, exceeding six months (Kohler et al., 2011). Thus, an empirical question remains, as it is unclear if these gene patterns represent a maintained state, or a transitory event. If they are time-limited events, the chance of registering such an event in the slice of time caught by a brain slice will be radically different in short- and long-lived mammals, and comparisons must take these basic scaling features into account.

## Acknowledgements

RNA sequencing data from human hippocampi were taken from the Allen Institute Website, which are supported by the NIH Contract HHSN-271-2008-00047-C to the Allen Institute for Brain Science. The opinions in this article are not necessarily those of the NIH. We thank Dr. Marnin Wolfe for helpful statistical discussions.

## Author Contributions Statement

BF and CC designed the study, analyzed the data, and wrote the study. All authors read and approved the manuscript.

## References

Andreae, L. C. (2018). Adult neurogenesis in humans: Dogma overturned, again and again?. Science Translational Medicine 10, eaat3893.

Albuixech-Crespo, B., López-Blanch, L., Burguera, D., Maeso, I., Sánchez-Arrones, L., Moreno-Bravo, J. A., et al. (2017). Molecular regionalization of the developing amphioxus neural tube challenges major partitions of the vertebrate brain. PLoS biology 15, e2001573.

Amrein, I., Nosswitz, M., Slomianka, L., van Dijk, R. M., Engler, S., Klaus, F., et al. (2015). Septo-temporal distribution and lineage progression of hippocampal neurogenesis in a primate *(Callithrix jacchus*) in comparison to mice. Front. Neuroanat. 9, 85.

Amrein, I., Isler, K., and Lipp, H. P. (2011). Comparing adult hippocampal neurogenesis in mammalian species and orders: influence of chronological age and life history stage. Eur. J. Neurosci. 34, 978–987.

Barton, R. A., and Venditti, C. (2013). Human frontal lobes are not relatively large. Proc. Natl. Acad. Sci. U. S. A. 110, 9001–9006. doi: 10.1073/pnas.1215723110

Ben Abdallah, N. M., Slomianka, L., Vyssotski, A. L., and Lipp, H. P. (2010). Early age-related changes in adult hippocampal neurogenesis in C57 mice. Neurobiol. Aging 31, 151–161.

Boldrini, M., Fulmore, C. A., Tartt, A. N., Simeon, L. R., Pavlova, I., Poposka, V., et al. (2018). Human hippocampal neurogenesis persists throughout aging. Cell Stem Cell. 22, 589–599.

Cahalane, D., Charvet, C.J., and Finlay, B.L. (2014). Modeling local and cross-species neuron number variations in the cerebral cortex as arising from a common mechanism. Proc. Natl. Acad. Sci. U.S.A. 111, 17642–17647. https://doi.org/10.1073/pnas.1409271111

Chaplin, T. A., H-H. Yu, J.G., Soares, Gatass, R., and Rosa, M.G.P. (2013). A conserved pattern of differential expansion of cortical areas in simian primates. J. Neurosci. 33, 15120–15125. doi: 10.1523/jneurosci.2909-13.2013

Charvet, C.J., Cahalane, D.J., and Finlay, B. L. (2015) Systematic, cross-cortex variation in neuron numbers in rodents and primates. Cereb. Cortex 25, 147–160. doi:10.1093/cercor/bht214

Charvet, C.J., Darlington, R.B., and Finlay, B. L. (2013) Variation in human brains may facilitate evolutionary change toward a limited range of phenotypes. Brain Behav. Evol. 81, 74–85 doi:10.1159/000345940

Charvet, C.J., and Finlay, B.L (2014). Evo-devo and the primate isocortex: the central organizing role of intrinsic gradients of neurogenesis. Brain, Behav. Evol. 84, 81–92 doi: 10.1159/000365181

Charvet, C. J., Hof, P. R., Raghanti, M. A., Kouwe, A. J., Sherwood, C. C., and Takahashi, E. (2017a). Combining diffusion magnetic resonance tractography with stereology highlights increased cross-cortical integration in primates. J. Comp. Neurol. 525, 1075–1093. https://doi.org/10.1002/cne.24115

Charvet, C. J., Šimić, G., Kostović, I., Knezović, V., Vukšić, M., Leko, M. B., et al. (2017b). Coevolution in the timing of GABAergic and pyramidal neuron maturation in primates. Proc. R. Soc. B. 284, 20171169. http://dx.doi.org/10.1098/rspb.2017.1169.

Charvet, C. J., and Striedter, G. F. (2008). Developmental species differences in brain cell cycle rates between northern bobwhite quail (*Colinus virginianus*) and parakeets (*Melopsittacus undulatus*): implications for mosaic brain evolution. Brain Behav. Evol. 72, 295–306.

Charvet, C. J., and Striedter, G. F. (2010). Bigger brains cycle faster before neurogenesis begins: a comparison of brain development between chickens and bobwhite quail. Proc. R. Soc. B. 277, 3469–3475.

Cipriani, S., Ferrer, I., Aronica, E., Kovacs, G. G., Verney, C., Nardelli, J., et al. (2018). Hippocampal Radial Glial Subtypes and Their Neurogenic Potential in Human Fetuses and Healthy and Alzheimer’s Disease Adults. Cereb. Cortex 28, 2458–2478.

Clancy, B., Darlington, R.B., and Finlay, B.L. (2000). The course of human events: predicting the timing of primate neural development. Dev. Sci. 3, 57–66

Clancy, B., Darlington, R.B., and Finlay, B.L. (2001). Translating developmental time across mammalian species. Neurosci 105, 7–17 doi.org/10.1016/S0306-4522(01)00171-3

Clancy, B., Kersh, B, Hyde, J., Anand, K.J.S, Darlington, R.B., and Finlay, B.L. (2007). Web-based method for translating neurodevelopment from laboratory species to humans. Neuroinformat. 5, 79–94 doi.org/10.1385/NI:5:1:79

Darlington, R.B., Dunlop, S.A., and Finlay, B.L. (1999). Neural development in metatherian and eutherian mammals: variation and constraint. J. Comp. Neurol. 411, 359–368

Dennis, C. V., Suh, L. S., Rodriguez, M. L., Kril, J. J., and Sutherland, G. T. (2016). Human adult neurogenesis across the ages: an immunohistochemical study. Neuropathol Appl. Neurobiol. 42, 621–638.

Dyer, M.A., Martins, R., da Silva Filho, M., Muniz, J.A., Silveira, L.C.L., Cepko, C., and Finlay, B.L. (2009). Developmental sources of conservation and variation in the evolution of the primate eye. Proc. Natl. Acad. Sci. U. S. A. 106, 8963 – 8968.

Fan, X., Dong, J., Zhong, S., Wei, Y., Wu, Q., Yan, L., et al. (2018). Spatial transcriptomic survey of human embryonic cerebral cortex by single-cell RNA-seq analysis. Cell research 28, 730–745.

Finlay, B.L., and Clancy, B. (2008) “Chronology of the development of the mouse visual system.” Eye, Retina and Visual System of the Mouse L. Chalupa and R.W. Williams, eds. (MIT Press: Cambridge, MA). pp 257–265

Finlay, B.L., and Darlington, R.B. (1995). Linked regularities in the development and evolution of mammalian brains. Science 268, 1578–1584 doi: 10.1126/science.7777856

Finlay, B.L., Darlington, R.D. and Nicastro, N. (2001). Developmental structure of brain evolution. Behav. Brain Sci. 24, 263–308 doi:10.1017/S0140525X01003958

Finlay, B.L., Hersman, M.N., and Darlington, R.B. (1998). Patterns of vertebrate neurogenesis and the paths of vertebrate evolution. Brain Behav. Evol. 52, 232– 242

Finlay, B.L., Hinz, F., and Darlington R.B. (2011). Mapping behavioral evolution onto brain evolution: The strategic roles of conserved organization in individuals and species. Phil. Trans. Roy. Soc. B. 366, 2111–2123 doi:10.1098/rstb.2010.0344

Finlay, B.L. and Workman, A.J. (2013). Human exceptionalism. Trends Cogn. Sci. 17, 199–201 doi: 10.1016/j.tics.2013.03.00

Finlay, B.L. and Uchiyama, R. (2017). “The timing of brain maturation, early experience, and the human social niche.” In: Kaas, J (ed.), Evolution of Nervous Systems 2e. vol. 3, pp. 123–148. Oxford: Elsevier.

Fleagle, J. G. (1984). ‘Size and adaptation in primates.’ The Lesser Apes: Evolutionary and Behavioral Biology. H. Preuschoft, D. Chivers, W. Brockelman and N. Creel. Edinburgh, Edinburgh University Press: 1–19.

Freckleton, R.P., Harvey, P.H., and Pagel, M. (2002). Phylogeneticanalysis and comparative data: atest and review of evidence. Am. Nat. 160, 712–726.

Garwicz, M., Christiansen, M., and Psouni, E. (2009). A unifying model for timing of walking onset in humans and other mammals. Proc.Natl. Acad. Sci. U. S. A. 106, 21889–21893. https://doi.org/10.1073/pnas.0905777106

Gould, S. J. (1975). “Allometry in primates, with emphasis on scaling and the evolution of the brain.” Approaches to Primate Paleobiology. Szalay. Basel, Karger. 5, 244–292.

Habib, N., Avraham-Davidi, I., Basu, A., Burks, T., Shekhar, K., Hofree, M., et al. (2017). Massively parallel single-nucleus RNA-seq with DroNc-seq. Nature methods 14, 955.

Halley, A. C. (2016). Prenatal brain-body allometry in mammals. Brain Behav. Evol. 88, 14–24. https://doi.org/10.1159/000447254

Halley, A. C. (2017). Minimal variation in eutherian brain growth rates during fetal neurogenesis. Proc. Roy.Soc. B: 284, pii: 20170219. DOI: 10.1098/rspb.2017.0219

Hawkes, K. and Finlay, B.L. (2018). Mammalian brain development and the evolution of our grandmothering life history. Physiol. Beh. 193, 55–68. https://doi.org/10.1016/j.physbeh.2018.01.013

Herculano-Houzel S., Collins C.E., Wong P., and Kaas, J.H. (2007) Cellular scaling rules for primate brains. Proc. Natl. Acad. Sci. U.S.A. 104, 3562–3567 doi: 10.1073/pnas.0611396104

Hikishima, K., Sawada, K., Murayama, A. Y., Komaki, Y., Kawai, K., Sato, N., et al. (2013). Atlas of the developing brain of the marmoset monkey constructed using magnetic resonance histology. Neurosci. 230, 102–113.

Hochgerner, H., Zeisel, A., Lönnerberg, P., and Linnarsson, S. (2018). Conserved properties of dentate gyrus neurogenesis across postnatal development revealed by single-cell RNA sequencing. Nat. Neurosci. 21, 290–299.

Hofman, M. A. (1989). On the evolution and geometry of the brain in mammals. Prog. Neurobiol. 32, 137–158.

Iacono, G., Benevento, M., Dubos, A., Herault, Y., Bokhoven, H., Kasri, N.N. and Stunnenberg, H.G. (2017). Integrated transcriptional analysis unveils the dynamics of cellular differentiation in the developing mouse hippocampus. Sci Rep 7, 18073.

Jabès, A., Lavenex, P. B., Amaral, D. G., and Lavenex, P. (2010). Quantitative analysis of postnatal neurogenesis and neuron number in the macaque monkey dentate gyrus. Eur. J. Neurosci. 31, 273–285.

Jerison, H. J. (1973). Evolution of the Brain and Intelligence. New York, Academic Press.

Jerison, H. J. (1989). Brain size and the evolution of mind. The 59th James Arthur Lecture on the Evolution of the Human Brain 1991: 1–99.

Jerison, H. J. (1997). “Evolution of prefrontal cortex.” Development of the Prefrontal Cortex: Evolution, Neurobiolgy and Behavior. N. A. Krasnegor, G. R. Lyon and P. S. Goldman-Rakic. Baltimore, Pall H. Brooks Publishing Co.: 9–26.

Kaas J.H., and Herculano-Houzel, S. (2017) “What Makes the Human Brain Special: Key Features of Brain and Neocortex”. In: Opris I., Casanova M. (eds) The Physics of the Mind and Brain Disorders. Springer Series in Cognitive and Neural Systems, vol 11. Springer, Cham https://doi.org/10.1007/978-3-319-29674-6_1

Kaushal, D., Contos, J. J., Treuner, K., Yang, A. H., Kingsbury, M. A., Rehen, S. K. et al. (2003). Alteration of gene expression by chromosome loss in the postnatal mouse brain. J. Neurosci. 23, 5599–5606.

Kempermann, G., Gage, F. H., Aigner, L., Song, H., Curtis, M. A., Thuret, S., et al. (2018). Human adult neurogenesis: evidence and remaining questions. Cell stem cell 23, 25–30

Knoth, R., Singec, I., Ditter, M., Pantazis, G., Capetian, P., Meyer, R. P., et al. (2010). Murine features of neurogenesis in the human hippocampus across the lifespan from 0 to 100 years. PloS one, 5, e8809.

Kohler, S. J., Williams, N. I., Stanton, G. B., Cameron, J. L., and Greenough, W. T. (2011). Maturation time of new granule cells in the dentate gyrus of adult macaque monkeys exceeds six months. Proc. Natl. Acad. Sci, U. S. A. 10326–31.

Kornack, D. R. and Rakic, P. (1998). Changes in cell-cycle kinetics during the development and evolution of primate neocortex. Proc.Natl. Acad. Sci, USA 95, 1242–1246.

Lee, H., and Thuret, S. (2018). Adult Human Hippocampal Neurogenesis: Controversy and Evidence. Trends in Molecular Medicine. 24, 521–522.

Lumsden, A. and Krumlauf. R. (1996). Patterning the vertebrate neuraxis. Science 274, 1109– 1115.

Merrill, D. A., Karim, R., Darraq, M., Chiba, A. A., and Tuszynski, M. H. (2003). Hippocampal cell genesis does not correlate with spatial learning ability in aged rats. J Comp. Neurol., 459, 201–207.

Ming, G.-l. and Song, H. (2005). Adult neurogenesis in the mammalian central nervous system. Ann. Rev. Neurosci 28, 223–250. https://doi.org/10.1146/annurev.neuro.28.051804.101459

Ngwenya, L. B., Peters, A., and Rosene, D. L. (2006). Maturational sequence of newly generated neurons in the dentate gyrus of the young adult rhesus monkey J Comp. Neurol 498, 204–216.

Passingham, R. E. (1985). Rates of brain development in mammals including man. Brain Behav Evol 26, 167–175.

Passingham, R. E. and Smaers, J. B. (2014). Is the prefrontal cortex especially enlarged in the human brain allometric relations and remapping factors. Brain Behav. Evol. 84, 156–166. https://doi.org/10.1159/000365183

Piumatti, M., Palazzo, O., La Rosa, C., Crociara, P., Parolisi, R., Luzzati, F., et al. (2018). Non-newly generated, “immature” neurons in the sheep brain are not restricted to cerebral cortex. J. Neurosci. 38, 826–842.

Puelles, L., Harrison, M., Paxinos, G., and Watson, C. (2013). A developmental ontology for the mammalian brain based on the prosomeric model. TINS 36, 570–578. dx.doi.org/10.1016/j.tins.2013.06.004

Rao, M. S., Hattiangady, B., and Shetty, A. K. (2006). The window and mechanisms of major age-related decline in the production of new neurons within the dentate gyrus of the hippocampus. Aging Cell 5.6, 545–558.

Reep, R., Darlington, R.B. and Finlay, B.L. (2007) The limbic system in mammalian brain evolution. Brain, Behav. Evol. 70, 57–70

Remtulla, S. and Hallet P. E. (1985). A schematic eye for the mouse and comparisons with the rat. Vis. Res. 25, 21–31.

Semendeferi, K., A. Lu, N. Schenker and H. Damasio (2002). Humans and great apes share a large frontal cortex. Nat. Neurosci. 5, 272–276.

Schoenemann, P. T. (2006). “Evolution of the size and functional areas of the human brain.” Ann. Rev. Anthro. 35, 379–406.

Sherwood, C. C. and Smaers J. B. (2013). What’s the fuss over human frontal lobe evolution? Trends Cogn. Sci. 17, 432–433.

Sorrells, S. F., Paredes, M. F., Cebrian-Silla, A., Sandoval, K., Qi, D., Kelley, K. W., et al. (2018). Human hippocampal neurogenesis drops sharply in children to undetectable levels in adults. Nature 555, 377–381.

Smith CL, Blake JA, Kadin JA, Richardson JE, Bult CJ, and the Mouse Genome Database Group. (2018). Mouse Genome Database (MGD)-2018: knowledgebase for the laboratory mouse. Nucleic Acids Res. 46 (D1): D836–D842.

Takahashi, T., R.S. Nowakowski and V. S. Caviness (1994). Mode of cell proliferation in the developing mouse neocortex. Proc. Natl. Acad. Sci. U.S.A. 91, 375–379.

Workman, A. D., Charvet, C. J., Clancy, B., Darlington, R. B., and Finlay, B. L. (2013). Modeling transformations of neurodevelopmental sequences across mammalian species. J. Neurosci. 33, 7368–7383. https://doi.org/10.1523/JNEUROSCI.5746-12.2013

Yopak, K., Lisney, T., Collin, S.E., Montgomery, J., Darlington, R.B. and Finlay, B.L. (2010). Brain scaling from sharks to primates: A highly conserved vertebrate pattern. Proc. Natl. Acad. Sci. U.S.A. 107, 12946–12951.

Zhong, S., Zhang, S., Fan, X., Wu, Q., Yan, L., Dong, J., et al. (2018). A single-cell RNA-seq survey of the developmental landscape of the human prefrontal cortex. Nature 555, 524–528.

